# A fuzzy encounter complex precedes formation of the fully-engaged TIR1-Aux/IAA auxin co-receptor system

**DOI:** 10.1101/781922

**Authors:** Sigurd Ramans Harborough, Arnout P. Kalverda, Gary S. Thompson, Martin Kieffer, Martin Kubes, Mussa Quareshy, Veselina Uzunova, Justyna M. Prusinska, Ken-ichiro Hayashi, Richard Napier, Iain W. Manfield, Stefan Kepinski

**Affiliations:** Centre for Plant Sciences (S.R.H., M.K., S.K.) and Astbury Centre for Structural Molecular Biology (A.P.K., I.W.M.), Faculty of Biological Sciences, University of Leeds, Leeds LS2 9JT, U.K.; School of Biosciences, University of Kent, Canterbury CT2 7NJ, U.K.; School of Life Sciences, University of Warwick, Gibbet Hill Road, Coventry CV4 7AL, U.K.; Department of Biochemistry, Okayama University of Science, Okayama 700-0005, Japan

**Author notes:** Correspondence should be addressed to S.K. **Author contributions** S.R.H. performed and analysed the majority of the experiments, assisted by A.P.K., G.S.T., M.K.^1^, K.H. and I.W.M. The SPR assay of cvxIAA was performed by M.K.^3^. The TIR1 protein production was facilitated by M.Q., V.U., J.M.P., M.K.^3^ and R.N. All authors commented on the manuscript. S.R.H., A.P.K., R.N., I.W.M. and S.K. wrote the manuscript.

## Abstract

The plant hormone auxin regulates almost every aspect of plant development via the TIR1/AFB-auxin-Aux/IAA auxin co-receptor complex. Within this ternary complex, auxin acts as a molecular glue to promote the binding of Aux/IAA transcriptional repressor proteins to SCF^TIR1/AFB^ ubiquitin-ligase complexes, thereby catalysing their ubiquitin-mediated proteolysis. A conspicuous feature of the crystal structure of the complex is a rare *cis* W-P bond within the Aux/IAA degron motif. To study receptor complex assembly, we have used NMR to determine the solution structure of the amino-terminal half of the Aux/IAA protein AXR3/IAA17, including the degron, both in isolation and in complex with TIR1 and auxin. We show that this region of AXR3 is intrinsically-disordered with only limited elements of structure and yet the critical degron W-P bond occurs with an unusually high (1:1) ratio of *cis* to *trans* isomers. We show that assembly of the co-receptor complex involves both auxin-dependent and -independent interaction events in which the disorder of the Aux/IAA is retained. Further, using the synthetic auxin molecule cvxIAA and by analysing specific Aux/IAA conformers, we show that a subset of auxin-dependent binding events occur away from the base of the canonical auxin binding pocket in TIR1. Our results reveal the existence of a fuzzy, topologically-distinct ternary encounter complex and thus that auxin perception is not limited to sequential, independent binding of auxin and then Aux/IAA to TIR1.

## Introduction

Auxin is a central signalling molecule in plant biology with roles in both the patterning of developmental events and the regulation of cellular growth. Much of this capacity for control arises from its ability to alter programmes of gene expression. This is achieved through a remarkably short signal transduction pathway that sees auxin promote the destruction of the Aux/IAA co-receptor/co-repressor proteins by interacting directly with both the Aux/IAA and a member of the TIR1/AFB family of F-box protein auxin co-receptors (Supplementary Fig. 1). The TIR1/AFB F-box proteins are the substrate-selection components of a multi-protein SCF-type E3 ubiquitin-ligase called SCF^TIR1/AFB^. The formation of the TIR1/AFB-auxin-Aux/IAA co-receptor complex promotes the ubiquitination and consequent degradation of the Aux/IAA protein by detaining it in the vicinity of ubiquitin-conjugating enzymes that associate with the core catalytic components of SCF^TIR1/AFB^. In this way the Aux/IAA proteins become polyubiquitinated and so targeted for degradation in the 26S proteasome (reviewed^1–3^). The rapid removal of Aux/IAAs in response to increases in auxin concentration prompts the derepression of the genes to which they are targeted via their interaction with the AUXIN RESPONSE FACTOR (ARF) family of DNA-binding transcription factors (reviewed^4,5^) (Supplementary Fig. 1). In higher plants the Aux/IAA and ARF families have multiple members with both overlapping and unique functions, providing a system rich in potential for complex control of gene expression in different cellular and developmental contexts^2^.

Aux/IAA proteins consist of four conserved domains, with domain I (DI) being associated with the transcriptional co-repressor activity of the Aux/IAA^6–8^, and domains III and IV mediating interaction with ARF transcription factors^9,10^. The region of the protein required for interaction with TIR1 is located within domain II (DII) of the protein. Within DII, a 13 amino acid degron motif has been defined that is necessary and sufficient for auxin-enhanced and ubiquitin-dependent proteolysis^11–13^.

The crystal structure of the fully-docked TIR1/AFB-auxin-Aux/IAA co-receptor complex has dominated our understanding of auxin perception^14^. This representation of the complex shows the auxin molecule, indole-3-acetic acid (IAA), bound in a pocket on the co-receptor protein TIR1 (hereafter called the auxin binding pocket) and entombed by the Aux/IAA core GWPPV degron motif (Fig. 1a,b). Intriguingly, the degron in this bound state shows a *cis*-prolyl imide bond between W86 and P87, raising questions about the impact of prolyl *cis-trans* isomerisation on the formation of the auxin co-receptor complex.

**Figure 1.**
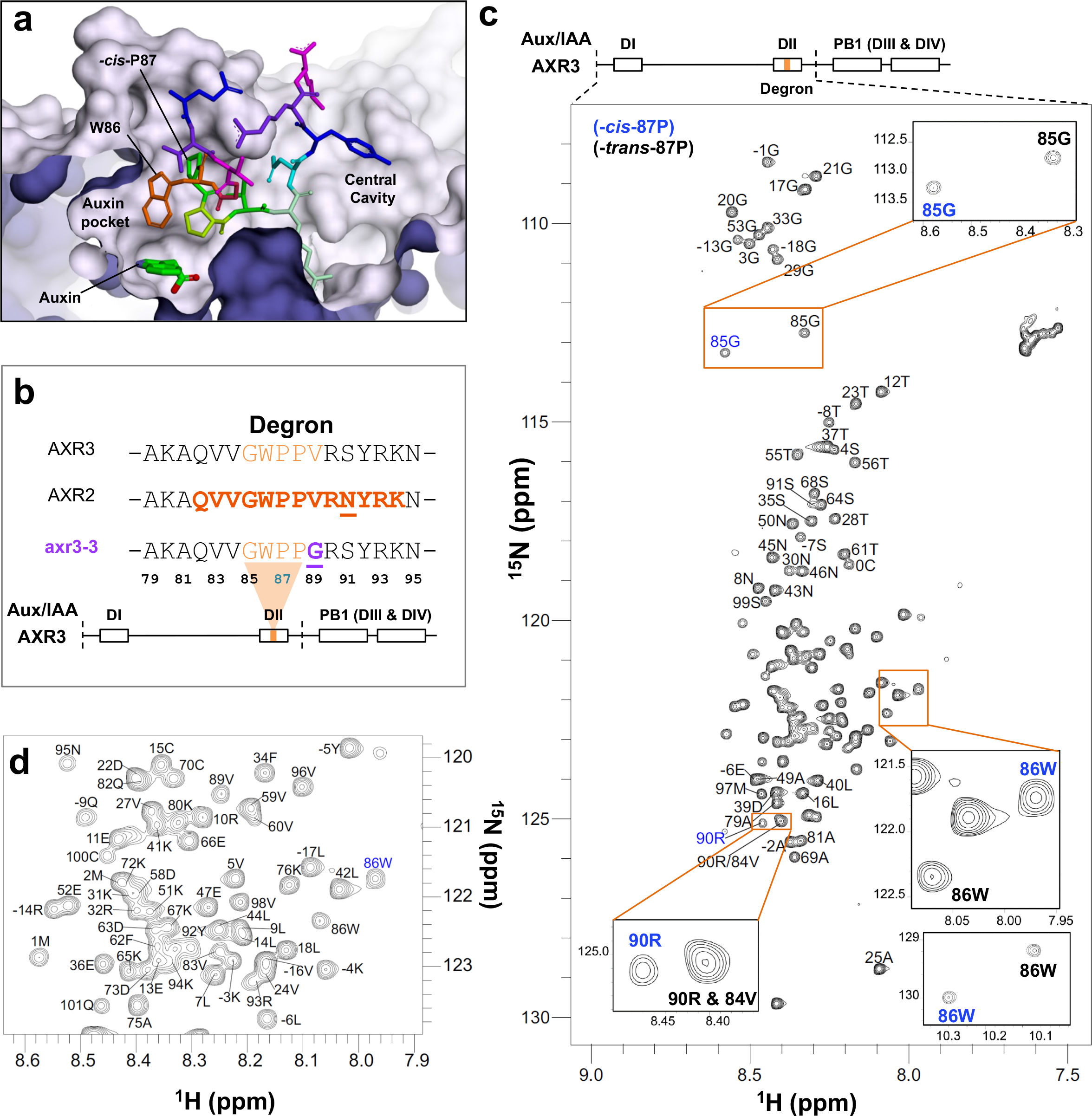
Overview of the Aux/IAA degron and the intrinsic disorder of AXR3 DI/DII. (**a**) Structure of IAA7/AXR2 degron (*cis*-P87) bound to TIR1 and auxin, showing the two TIR1 cavities based on 2P1Q (Tan *et al.*, 2007). (**b**) Amino acid sequences of DII from different Aux/IAA proteins with polymorphisms highlighted in bold and underlined. Core residues are in orange, and the mutated residue in axr3-3 is shown in purple. The AXR2 sequence highlighted and in bold indicates the peptide crystallised by Tan *et al.*, 2007^14^. Below the sequence alignment is a schematic of the AXR3 protein showing the four domains. The location of the degron is highlighted, and the dashed line indicates the DI/DII region of the protein studied by NMR (**c**) ^1^H-^15^N HSQC spectrum of the protein AXR3 DI/DII at 16.5 °C. The peaks associated with P87 in the *cis* isomer conformation are annotated light-blue. (**d**) An enlarged image of the signal dense region of the HSQC spectrum.

Indeed, specific sequence contexts in the Aux/IAA degron may establish the inherent stability of the *cis*-P87 isomer state. At a basal level, the typical population for a *cis* imide bond in a protein structure is 5-6%^15^. The *cis* population is enhanced when an aromatic amino acid precedes the prolyl bond^16^. From analysis of short peptides, the sequence combination W-P is expected to result in a 25% *cis* population for the imide bond between the two residues, with a decrease in the isomerisation rate from *cis* back to *trans*^17^. This *cis* isomer population is expected to be further enhanced by an additional C-terminal proline residue, giving a di-proline motif (in the AXR3 degron W86-*cis-*P87*-trans-*P88), where the *cis-trans* conformer population is expected to be 37%^18^, a value very close to the 36% previously reported for the degron W-P bond in peptides for the rice Aux/IAA OsIAA11^19,20^.

To date there has been no structural information of a full-length Aux/IAA protein. The most complete structures of Aux/IAA proteins are of the carboxy-terminal (C-terminal) half of the protein including domains DIII and DIV/PB1^3,10^, which are not directly involved in auxin perception. The amino-terminal (N-terminal) half (DI and DII) of the Aux/IAA, including the degron motif that is required for interaction with TIR1/AFBs and hence auxin sensing, is predicted by bioinformatics to be intrinsically disordered^21^. In broad terms, intrinsic disorder describes a protein or protein region that cannot be fully represented by a single 3D structure^22,23^. Intrinsically disorder in proteins can be further refined into classifications of static or dynamic disorder^22^. Static disorder refers to a protein or protein region that adopts multiple stable conformations. Where a protein constantly fluctuates between conformations the disorder is said to be dynamic. These distinctions can be extended to protein-protein complexes involving disordered proteins, which are termed as being fuzzy^22^. Such complexes can show static fuzziness, where disordered regions become ordered upon binding but form more than one conformation^22,23^. Where the disordered protein or protein region fails to undergo a disorder-to-order transition upon binding the fuzziness of the complex is said to be dynamic^22^. The existence, nature, and fate of any instrinsic disorder within Aux/IAAs has until now been unknown.

Here, we have used nuclear magnetic resonance (NMR) spectroscopy to characterise the solution structure of the 101 amino acid N-terminal half (DI/DII) of the Aux/IAA protein AXR3 both in isolation and in complex with TIR1 and auxin. We show that the N-terminal half of AXR3 is intrinsically disordered and yet contains elements of transient secondary structure and an unusually high occupancy of the *cis* state for a critical W-P bond within the degron motif. This W-*cis*-P degron conformer supports the highest level of ternary complex formation, assisted by a binding interface which extends C-terminal to the degron. Throughout receptor complex formation, the disordered state of the Aux/IAA remains. We show that there are stages of complex assembly that are mediated by auxin-dependent and -independent binding events occurring away from the base of the auxin binding pocket in TIR1. In addition to indicating the existence of an early, lower-affinity encounter complex these data demonstrate that receptor complex assembly is not limited to the sequential binding of auxin and then Aux/IAA to TIR1, revealing additional steps at which functional specificity in the auxin perception process could arise. Together these data provide a framework for understanding the process of auxin perception through the formation of a fuzzy auxin receptor complex.

## Results

### Structural characterisation of AXR3 domains I and II (DI/DII)

We used solution-state NMR spectroscopy to study the structure of DI and DII of AXR3 (AXR3 DI/DII, residues 1 to 101, Fig. 1b). We specifically chose to exclude the C-terminal PB1 domain to avoid multimerisation that would otherwise obfuscate the analysis of the regions of the protein directly involved in auxin perception. The N-terminal half (DI/DII) of AXR3 and the mutated variant axr3-3 DI/DII were expressed as ^13^C, ^15^N isotopically-labelled proteins (Fig. 1b, methods: protein preparation). The following NMR backbone assignment experiments were performed: HNCA, HNcoCA, HNcaCB, CBcacoNH, HNcaCO, HNCO, CON, hCACO, hCAnCO (Supplementary Tables 1 and 2). Together, these experiments resulted in the assignment of 99% of resonances in AXR3 DI/DII and the variant proteins.

Our data show that the AXR3 degron is situated in an extensive region of dynamic intrinsic disorder, encompassing the majority of domains I and II. The ^1^H-^15^N Heteronuclear Single-Quantum Correlation (HSQC) spectrum for AXR3 DI/DII is characterised by signals occurring in a narrow ^1^H chemical shift region (7.9 to 8.6 ppm) (Fig. 1c,d). Despite extensive intrinsic disorder, our analyses of chemical shift indices and ^15^N R_2_ relaxation rates also show a propensity to form secondary structure, which is transiently formed and observed only in a subset of the protein population at any one time^24^. This includes helical character at the N-terminus and a preference for extended structures in the β-region of phi/psi space at the C-terminus (Fig. 2a, Supplementary Figs. 2 & 3).

**Figure 2.**
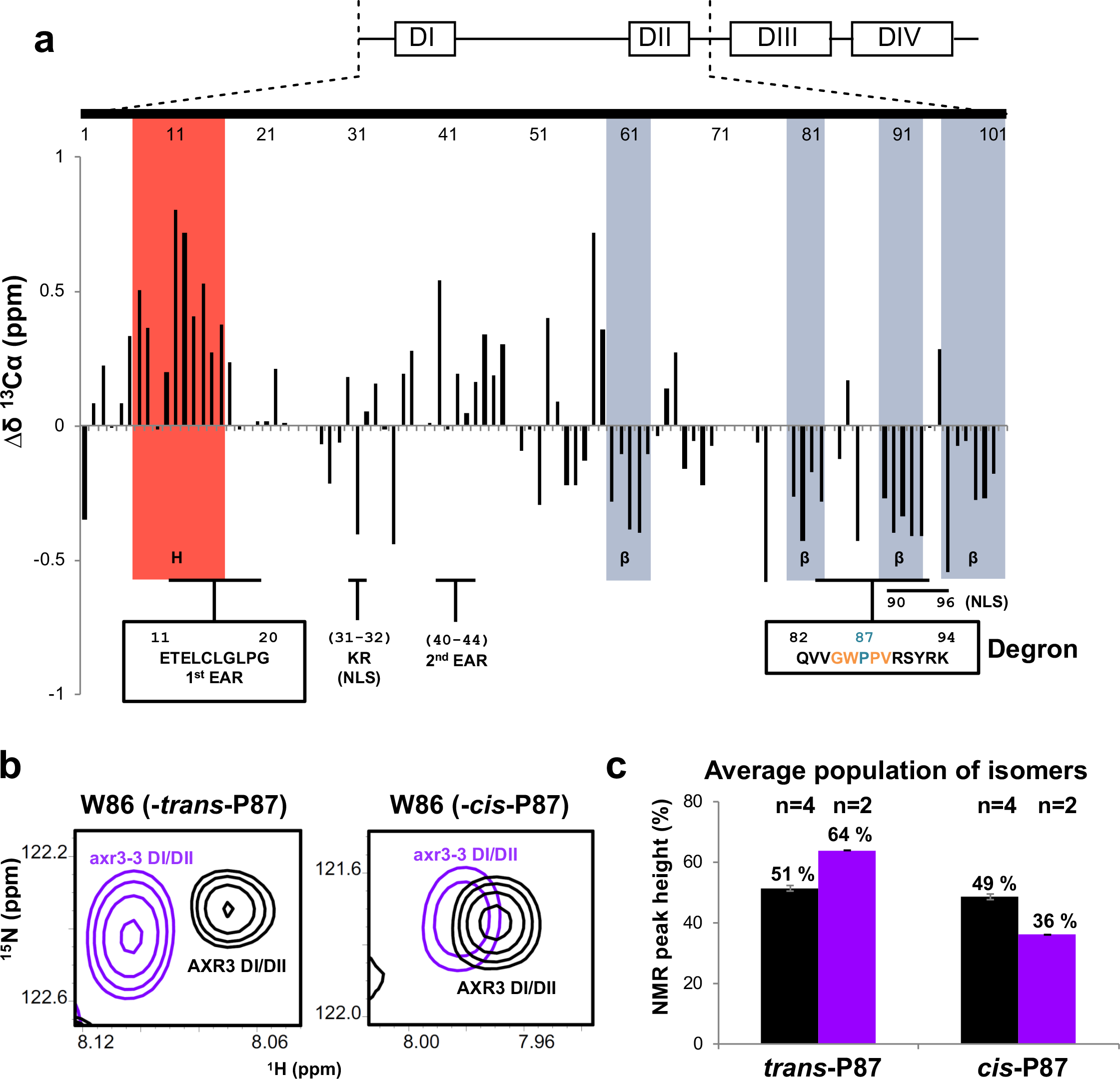
Proline 87 within the degron core of AXR3 exhibits a high *cis*:*trans* ratio despite the lack of substantial stable structure in the vicinity of the degron. (**a**) The chemical shift index for ^13^C_α_ signals assigned to residues along the carbon backbone of AXR3 DI/DII. Positive chemical shift differences (Δδ) indicate a tendency for helical secondary structure. Negative Δδ indicate a tendency for β-secondary structure. Consensus regions for secondary structure tendencies between the Δδ, ^13^C_α_, and ^13^C’ indexes are highlighted red and grey, for the helical and β-secondary structure respectively (Supplementary Fig. 2). The data are overlaid with a schematic diagram of the AXR3 protein, indicating the N-terminal DI and DII. Important binding interfaces include the EAR motifs for the recruitment of TPL. The two nuclear localisation signals (NLS) are also indicated. (**b**) HN cross peaks associated with W86 in AXR3 DI/DII compared to axr3-3 DI/DII. (**c**) The *cis* and *trans* isomer populations determined from the HN cross peak heights recorded for the G85 and W86 signals in axr3-3 DI/DII and AXR3 DI/DII. In each study the mean is calculated and n = number of independent studies. The overall mean of means is shown and the error bars display the standard deviation.

The nascent helical region corresponds to residues 7-16 of DI and includes the majority of the primary EAR motif of LXLXL (Fig. 2a and Supplementary Fig. 2), which forms a key interface for the recruitment of the co-repressor TOPLESS (TPL) (Supplementary Fig. 1). In contrast, the degron is situated (residues 82 to 94) within a cluster of weak β-secondary structure (Fig. 2a and Supplementary Fig. 2). In addition to these regions with propensity for secondary structure, ^15^N R_2_ relaxation rates show a more complex profile than would be expected for a completely disordered chain, indicating other long-range interactions or hydrophobic clustering are likely to be present (Supplementary Fig. 3).

### Proline 87 within the degron core of AXR3 exhibits a *cis* and *trans* ratio of 1:1

Within the degron core (residues 85 to 89) the prolyl bond between W86 and P87 is of particular interest because of its *cis* conformation in the crystal structure of the co-receptor complex^14^ (Fig. 1a). NMR analysis of the AXR3 DI/DII protein shows clear splitting of resonances for degron residues V84 to R90 arising from the *cis* and *trans* isomers of P87, but the *cis* linked-states for resonances V84 and V89 are obscured by the overlap of peaks in the ^1^H-^15^N HSQC spectrum (Fig. 1c and 1d). This also affected the *trans*-linked resonances for V84 and R90, which were only distinguished at 4°C and 950 MHz. Therefore, the populations of the isomer states were determined from the height of the HN cross peaks associated with G85 and W86 and found to be 49:51, *cis* and *trans* at 16.5°C (Fig. 1c, Fig. 2b and 2c). Even for an X-Pro imide bond with a predicted ratio of 37:63 in a random coil, this represents a remarkably high proportion of the *cis* isomer, particularly in the context of the largely disordered nature of this region of the protein.

To assess the contribution of closely adjacent degron residues to P87 isomerisation state, we performed NMR analysis of the N-terminal half of AXR3 bearing the axr3-3 mutation (axr3-3 DI/DII), where V89 is changed to glycine (WPPV to WPPG)^25,26^. This mutation is part of the *axr3* mutant series that has been well characterised at the whole plant level. Mutant phenotypes include reductions in plant stature and gravitropic response, and severity is associated with proximity of the mutated residue to the degron W-P bond, with *axr3-1* (P88L) being more severe than *axr3-3* (V89G)^25,26^. Consistent with the dominant, gain-of-function nature of the phenotypes, the axr3-1 mutation has been shown to abolish interaction with SCF^TIR1/AFB^, leading to stabilisation and over-accumulation of the protein^11,27^. ^1^H-^15^N HSQC analysis of the axr3-3 DI/DII protein shows that the valine to glycine substitution at position 89 shifts the *cis* and *trans* ratio to approximately 1:2, decreasing the amount of the *cis-*P87 state (Fig. 2b and 2c). These data demonstrate that despite the lack of substantial stable structure in the vicinity of the degron, adjacent residues contribute to the *cis-trans* isomerisation of the W86-P87 imide bond.

### Auxin-enhanced binding of the *cis*-P87 conformation to TIR1

The narrow topography of the auxin pocket in TIR1 is expected to dramatically restrict accommodation of the *trans* conformer, which is not observed in the crystal structure^14^. To study the impact of *cis* and *trans* degron isomers on auxin co-receptor complex formation we performed ^1^H-^15^N HSQC NMR. AXR3 DI/DII protein, isotopically labelled with ^15^N, was assayed with unlabelled TIR1 in the presence or absence of the unlabelled auxin (IAA) (Fig. 3a). Changes in ^15^N/^1^H chemical shift, and/or decreases in signal intensity due to line-broadening are common signs of a molecular interaction in an NMR experiment^28^. Especially in the case of interactions between an intrinsically disordered protein and a folded protein, signal intensity decreases and disappearance of signals rather than chemical shift changes are often observed as a result of the interaction^29^. The degree of line-broadening is dependent on a number of factors. These include the exchange rate between the free and receptor bound state, the magnitude of any chemical shift, and the difference in the intrinsic peak line width between free and bound state (T_2_). Therefore, for both very strong and very weak interactions the degree of line broadening is limited, due to the exchange rate being too slow or too fast respectively. In addition, signal intensity decreases can still be seen outside of the direct interaction surface as a result of a difference in rotational correlation time between free and receptor bound protein (Fig. 4a). Consequently, for peaks which were well resolved and not overlapping within the spectra, decreases in the NMR signal intensity were used as an indicator of binding.

**Figure 3.**
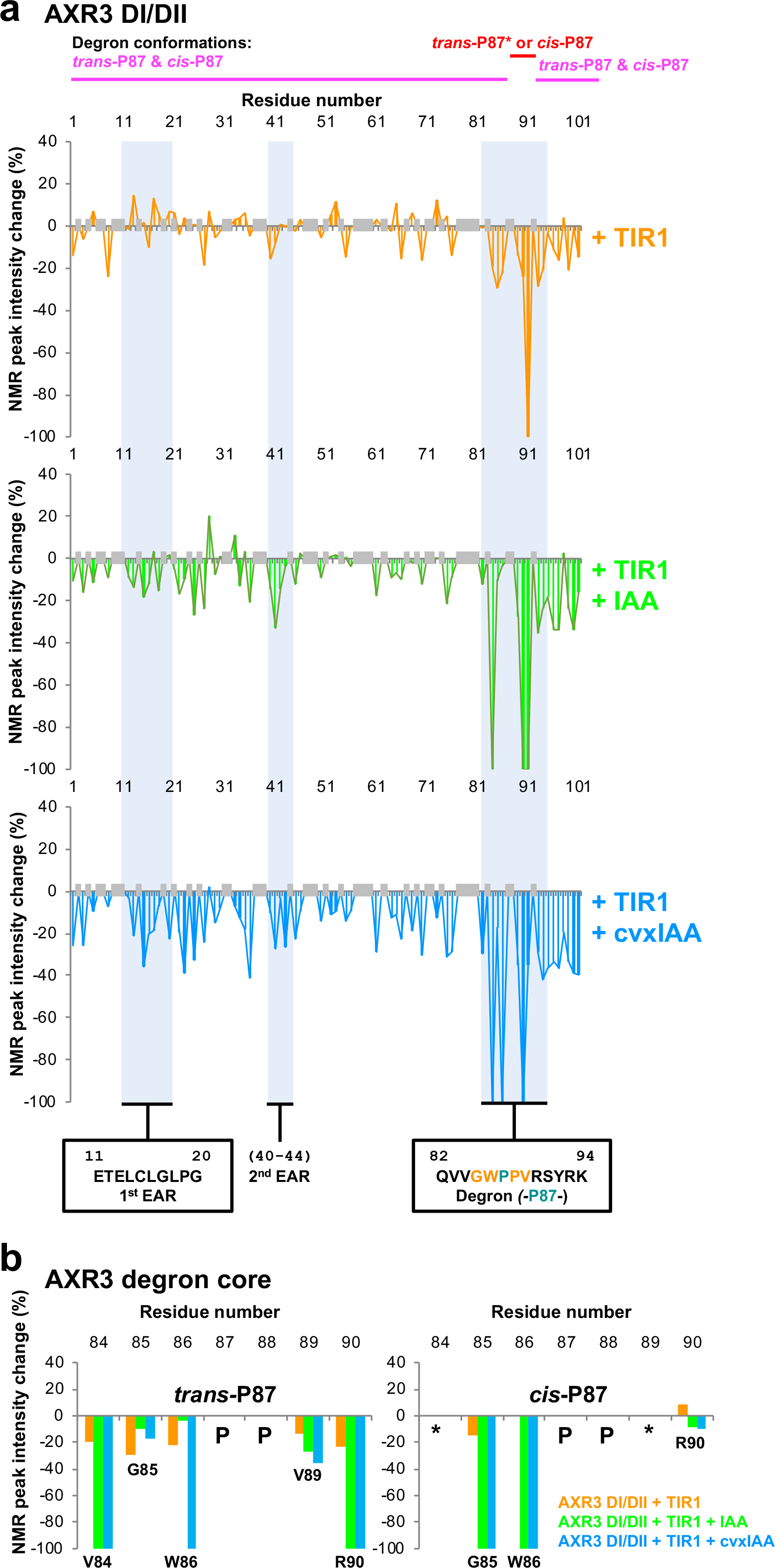
Co-receptor complex formation involves both auxin-dependent and -independent events, including ternary complex formation away from the base of the auxin binding pocket in TIR1. (**a**) Percentage changes in the intensity of HN cross peaks from ^1^H-^15^N HSQC spectra of AXR3 DI/DII with the addition of TIR1 (orange), and TIR1 with IAA (green). A change of −100% indicates that peak intensity has decreased to the noise floor and is no longer observed. Important binding interfaces are annotated and include the EAR motifs and the degron, these regions are shaded blue on the graphs. Degron conformation labels at the top of the figure highlight regions common to both *cis*- and *trans*-P87 degron conformations in magenta and the region V84 to R90 across which clear splitting of resonances associated with either *cis*- or *trans*-P87 degron conformers in red (*percentage changes for the *trans*-P87 degron are shown in (**a**), with data for both *trans*-P87 and *cis*-P87 V84 to R90 in (**b**)). Grey shaded bars on the graph indicate residues for which peak intensity could not be measured due to HN peak overlap or where prolines are positioned in the sequence. (**b**) Percentage differences for the AXR3 degron with the addition of TIR1 and IAA. Missing data points where the peak intensity could not be measured due to HN peak overlap are indicated with the symbol (*); prolines are indicated with the symbol (P).

**Figure 4.**
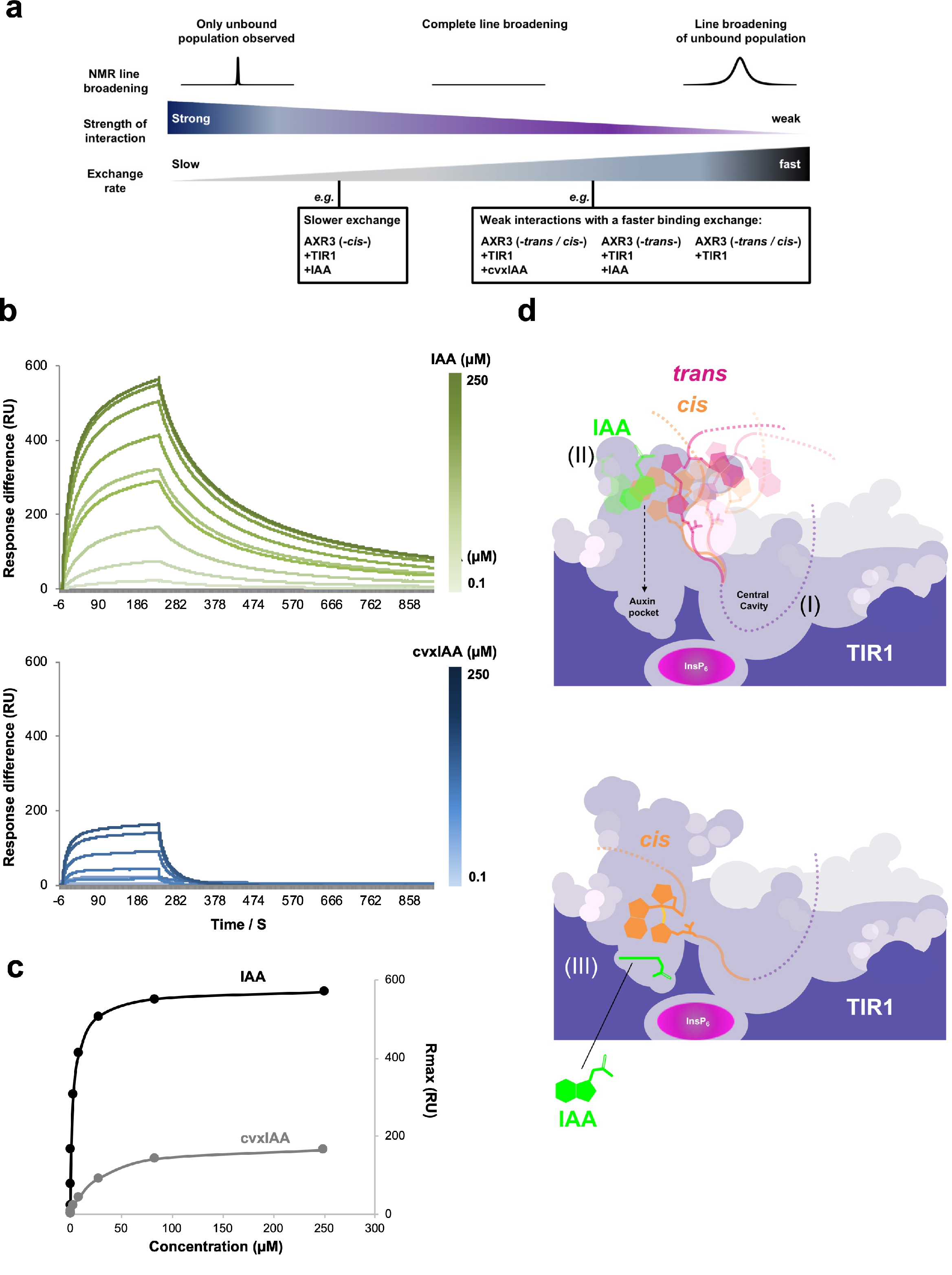
Affinity and assembly of IAA- and cvxIAA-based auxin co-receptor complexes. (**a**) The relationship between NMR peak intensity and binding kinetics. The interaction of AXR3 and TIR1 with and without auxin was investigated by NMR. In these assays it is the AXR3 protein which is labelled and the focus of the observation. The interaction is observed through the association and dissociation of AXR3 from the receptor, where a binding event is represented by decreases in the NMR peak intensities and the magnitude of this change is dependent on the binding kinetics of the interaction. In NMR there is an optimum of binding kinetics for complete line-broadening, and either side of this optimum the peak intensities of this protein population will start to increase. Therefore, the magnitude of NMR peak intensity change will vary according to the affinity and binding exchange rate of the interaction being studied as indicated. (**b**) SPR analysis of the kinetics of binding between AXR2 degron peptide and TIR1 induced by IAA or cvxIAA (upper and lower panel respectively) (**c**) The response amplitude of sensorgram data shown in (b) plotted against auxin concentration. (**d**) Schematic representation of a cross section through the TIR1 receptor showing events during the assembly of the auxin co-receptor complex. (I) Auxin-independent binding of the Aux/IAA, particularly C-terminal to the degron core. (II) Encounter complex, where auxin (IAA) promotes interaction of the degron core and TIR1 away from the base of the auxin binding pocket. The *cis* and *trans* conformers of the degron are shown in orange and pink respectively. The transparency of the representations of the degron conformers and IAA indicate uncertainty over precise positioning during this encounter phase. (III) The lower panel shows the final binding pose of the degron based on the crystal structure^14^. The Aux/IAA degron in the *cis*-conformer is fully engaged with TIR1, with auxin at the base of the pocket acting as molecular glue^14^.

Analysis of the spectra for AXR3 DI/DII protein alone and in complex with both TIR1 and auxin shows that the intrinsic disorder of AXR3 DI/DII, characterised by the congestion of signals in a narrow ^1^H chemical shift region, persists during the interaction (Supplementary Fig. 4) indicating that the ternary auxin co-receptor complex is a dynamic, fuzzy one^22^. Decreases in NMR peak intensity associated with the addition of TIR1 and IAA show the degron in DII is the primary binding interface, supported by the adjacent C-terminal region (Fig. 3a). The G85 and W86 HN cross peaks associated with the *cis* isomer of P87 display some of the largest changes when IAA is present, leading to the NMR signals no longer being observed due to broadening (Fig. 3b). It is notable that in the *trans* state of the degron, these two core residues only show limited engagement with TIR1 in the presence of auxin (Fig. 3b). These results indicate that *cis*-P87 locks the degron core onto the TIR1 surface and supports the strongest, auxin-enhanced binding to TIR1, consistent with earlier crystallographic data^14^. Further, our NMR analysis show that the binding interface extends beyond the peptide sequence used in these crystallographic studies (ending at K94)^14^. We observed changes in signal intensity to residue Q101 at the C-terminus of AXR3 DI/DII. Even at this distal location, auxin enhances AXR3 binding to TIR1 (Fig. 3a). Thus, the binding interface between the two proteins is likely to extend well into the central cavity of the solenoid of LRRs on the TIR1 surface (Fig. 1a; Tan *et al.*, 2007^14^), a finding that is consistent with previous studies in yeast showing that the region C-terminal to the degron in several Aux/IAAs is important for their instability^30^.

### Auxin-dependent binding of the *trans* degron to TIR1 and the synthetic auxin molecule cvxIAA reveal intermediate stages of co-receptor complex formation

In addition to the ternary co-receptor complex, we also observed decreases in NMR signal intensity in spectra obtained with only AXR3 DI/DII and TIR1. This is particularly evident in the C-terminal half of the *trans* degron (Fig. 3a, top and 3b). This region includes S91, where the signal intensity decreases to such an extent that the resonance is no longer detected, indicating the existence of auxin-independent association between AXR3 DI/DII and TIR1 (Fig. 3a).

Intriguingly, auxin-dependent binding between the *trans*-P87 degron conformer and TIR1 is detected for residues V84, V89, and R90 (Fig. 3b). These interactions are particularly interesting because the profound difference in the conformation of the *trans*-P87 degron and confines of the TIR1 auxin binding pocket preclude the canonical molecular glue mode of auxin activity^14^. These data therefore indicate formation of a ternary Aux/IAA-auxin-TIR1 complex away from the base of the pocket. Whether or not the auxin-induced binding of *trans*-P87 degron conformers to TIR1 is in itself of direct physiological relevance in auxin perception is unclear. One explanation for the observation is that the *trans*-P87 conformation, which cannot proceed to the fully-assembled complex defined by crystallographic analysis, reveals an auxin-dependent and topologically-distinct intermediate binding event common to both *trans* and *cis* degron conformations. In the case of the latter, this intermediate state is less readily detectable in the latter because of the rapid progression of *cis* degron conformers to the fully-docked state. To test this idea we performed ^1^H^−15^N HSQC NMR experiments using the auxin analogue 5-(3-methoxyphenyl)-indole-3-acetic acid (cvxIAA)^31^. This synthetic auxin, which is unable to dock at the base of the binding pocket in wild-type TIR1, only elicits auxin effects *in planta* in the presence of a TIR1 derivative modified to accommodate the methoxyphenyl side-group (ccvTIR1)^31^. cvxIAA thus provides a tool to study putative auxin-enhanced interactions occurring before the co-receptor complex is fully docked^32^.

Decreases in NMR peak intensity associated with the addition of TIR1 and cvxIAA were similar to the binding profile observed with IAA, but with larger changes to peak intensity (Fig. 3a). While a larger effect of cvxIAA relative to IAA may seem counter-intuitive, the more pronounced line-broadening with cvxIAA is consistent with the NMR behaviour of an interaction with an intermediately-fast binding exchange rate relative to the more stable interaction promoted by IAA (Fig. 4a). Indeed, Surface Plasmon Resonance (SPR) analysis of the receptor complex formation with cvxIAA shows much faster dissociation and an almost 10-fold decrease in affinity compared to IAA (Fig. 4b,c Supplementary Table 3). Importantly, cvxIAA promotes the binding of both conformers of the degron. This was particularly clear for the *cis* degron core, with the loss of signals for the HN cross peaks of G85 and W86, in contrast to the TIR1 only control (Fig. 3b). These results show that a binding event occurs between AXR3 DI/DII and TIR1, enhanced by cvxIAA, before the degron core is fully docked. These data, together with the observation of auxin (IAA)-dependent binding of *trans*-P87 AXR3 DI/DII to TIR1 indicate the existence of an encounter complex that, in terms of the spatial arrangement of the interacting co-receptor components, is topologically distinct from the fully-docked co-receptor complex observed by X-ray crystallography^14^ (Fig. 4d).

## Discussion

The data presented here have revealed the early events of auxin perception via TIR1/AFB and Aux/IAA co-receptor proteins. Central to this is the experimental demonstration of extensive intrinsic disorder in the DI/DII half of AXR3, consistent with previous bioinformatic predictions^21^. The local secondary structure propensity assignments for AXR3 reported here are supported by the report of α-helical structure for an IAA27 EAR motif peptide bound to the Arabidopsis TOPLESS protein^8^ (PDB code 5NQV) and helical configurations reported for peptides *At*IAA1 and *At*IAA10 peptides bound to rice TOPLESS-related protein 2^7^.

The intrinsically disordered region of AXR3 forms an auxin co-receptor complex primarily via the degron. This binding motif is surrounded by a cluster of weak β-secondary structure and the transient formation of such elements has previously been shown to have the potential to modulate the molecular interaction^33^. Together with the demonstration of auxin-independent and crucially, auxin-dependent AXR3-TIR1 binding events occurring away from the base of the auxin binding pocket in TIR1, we propose a model in which the C-terminal half of the degron and adjacent non-degron motifs initiate an encounter complex with its co-receptor, enhancing the likelihood of an active ternary complex being formed (Fig. 4d). The initial stages of this process are supported by evidence of auxin-independent association of the AXR3 proteins to TIR1, mediated predominantly through the C-terminal half of the degron and possibly driven by the electrostatic field of the InsP_6_ co-factor^14^. This evidence of auxin-independent association is consistent with previous pull-down assays using full-length Aux/IAA proteins^12,13^, native polyacrylamide gel electrophoresis^14^, and yeast two-hybrid assays^34^.

The auxin-independent association of Aux/IAA and TIR1 is likely to increase the probability of the degron core coming into close proximity to the entrance of the auxin binding pocket (Fig. 4d). In this configuration and with the inherent conformational flexibility of the region surrounding the degron, we speculate that the hydrophobic cluster of the degron core (VVGWPPV) may be primed for auxin to trigger the central degron to interact with TIR1, enhancing the transition of both auxin and the Aux/IAA degron further into the auxin binding pocket and into the positions as captured in the crystal structure^14^. The discovery of encounter complex interactions between TIR1, auxin, and AXR3 does not preclude less sophisticated modes of auxin perception in which auxin and the Aux/IAA co-receptor bind sequentially and independently to TIR1. Rather they add to the set of distinct molecular interaction events within which specificity in auxin signalling can arise^2,34,35^. Thus, the capacity for a small molecule to act as a potent auxin in promoting receptor complex formation is not limited to its ability to navigate to, and occupy the base of the auxin binding pocket in TIR1. Similarly, the intrinsic instability of an Aux/IAA is not dependent solely on its ability to occupy the space above auxin when bound at the bottom of that same pocket.

An intriguing observation of this work on AXR3 is that P87 shows approximately equal occupancy of the *cis* and *trans* conformations, without catalysis, observed through the splitting of the W86 and G85 HN cross peaks. This very high proportion of *cis* conformer is perhaps all the more remarkable given the paucity of substantial structure in the region of the degron. Nevertheless, the finding that mutation of the adjacent degron residue V89 dramatically reduces the *cis:trans* ratio from 1:1 to 1:2 indicates that the local environment of the degron contributes to the propensity to adopt the *cis* conformation. Previous work with degron peptides for the rice Aux/IAA, OsIAA11 reported a *cis* population of 36%^19^. That work identified a *cis-trans* isomerase, LRT2, which is proposed to be required to promote conversion between the two conformers^19, 20^. Recently, it has been shown that catalysis of bond of the OsIAA11 degron is intrinsically slow, regardless of which isomerases are used^36^. Indeed, the isomerisation rate without catalysis for a typical W-P bond is 5 ×10^−4^ s^1^ for *cis* to *trans*, and 3 ×10^−4^ s^−1^ for *trans* to *cis* at 4 °C^16^. With this in mind, no such isomerase has as yet been identified for Arabidopsis, but the possibility remains that there may be more to the function of TIR1 than previously thought.

By studying one of the most pivotal molecular interaction events in plant development, we have drawn attention to the interplay between plant hormones and intrinsically disordered proteins. These findings have identified new steps and new complexity in the formation of the Aux/IAA-TIR1 co-receptor complex involving at least two distinct auxin-dependent interaction events in addition to their auxin-independent association. Protein partners that retain elements of intrinsic disorder in one or more of their constituent parts upon complex formation, as in the case of AXR3 and TIR1, have been described as fuzzy^22^. In the case of TIR1-based auxin perception an interesting question arises as to whether the nanomolar affinity of the ternary co-receptor complex^12,13,37^ is remarkable despite the persistent intrinsic disorder of the Aux/IAA or because of it. In addition to providing the multivalency to support the formation of complexes involving multiple steps and binding surfaces, there is a growing body of work highlighting the crucial role that intrinsically disordered protein regions can play in tuning protein function^23,38,39,40^. For example, it has recently been shown that the intrinsically disordered tail of human UDP-α-d-glucose-6-dehydrogenase (UGDH) can alter the conformation and activity of the protein via the entropic force arising simply from constraint of an unfolded peptide against folded protein domains^39^. The entropic force generated was shown to be proportional to the length of the intrinsically disordered region and independent of amino acid sequence *per se*^39^. In the case of Aux/IAAs, it has been shown that truncations of AXR3/IAA17 that retain the sequences immediately C-terminal to the degron shown to interact with TIR1 (Fig. 3) but remove portions of the intrinsically disordered regions N-terminal to the degron increase protein half-life in an in vivo yeast system^30^. Further work will be required to establish the extent to which the inherent dynamic switching of conformational states of the intrinsically disordered regions of AXR3 contributes to either or both the high propensity for the *cis*-P87 degron form and the formation of Aux/IAA-auxin-TIR1 encounter complex and subsequent progression to the fully-docked co-receptor architecture. In addition, the new insights into the auxin co-receptor complex formation described here provide a framework to better understand the mode of action and species selectivity of current auxinic herbicides as well as highlighting the potential for the development of more selective and potent herbicides based on the regulation of a fuzzy auxin co-receptor complex.

## Methods

### Protein preparation

The N-terminal half DI/DII of AXR3, and axr3-3 were expressed as 6X His-tag (N-terminal) fusion proteins in *Escherichia coli* strain Rosetta™ DE3 competent cells (Supplementary Table 4; Novagen, product code: 70954). These proteins were expressed in minimal media with ^13^C D-glucose and ^15^N ammonium chloride. The maximisation of isotope labelling of the expressed protein involved a 125-fold dilution of cell culture in enriched growth media into minimal media with ^13^C D-glucose and ^15^N ammonium chloride and grown for 16 hours (37°C / 200 rpm); followed by a further 40-fold dilution into minimal media for the final period of cell growth and protein expression (induced with 0.5 mM IPTG / 18°C / 200 rpm and grown for a further 12 hours). The fusion protein was isolated from soluble cell lysate by Co-NTA affinity chromatography and the protein eluted on a gradient of increasing imidazole concentration. Chromatography buffers contained 20 mM sodium phosphate, pH 8.0, 500 mM NaCl and either 10 mM or 500 mM imidazole for wash and elution buffers respectively. For preparation of unlabelled Arabidopsis TIR1, expression constructs were engineered into the pOET5 transfer vector (Oxford Expression Technologies) to allow coexpression of the fusion proteins His10-eGFP-FLAG-(TEV)-AtTIR1 and His10-(TEV)-AtASK1 (pOET5 AtTIR1 AtASK1, Supplementary Fig. 5) in a baculovirus vector system in *Spodoptera frugiperda*9 (Sf9) insect cells and purified as previously described^34^ with the following modifications. Soluble cell lysate was passed through a HiTrap 1 mL TALON Crude column, followed by a column of ANTI-FLAG^®^ M2 affinity gel (Sigma-Aldrich, product code: A2220) with the sample in a buffer containing 1 mM DTT, 150 mM NaCl and 10 mM HEPES pH 7.4 and eluted with 100 μg ml^−1^ 3×FLAG peptide (Sigma).

### NMR spectroscopy

All protein samples for NMR analysis were concentrated by ultrafiltration and underwent buffer exchange into 20 mM sodium phosphate pH 6.0, 150 mM NaCl, 3 mM EDTA, 10 mM DTT, cOmplete mini protease inhibitor cocktail (2% v/v; Roche Molecular Biochemicals). Before NMR analysis D_2_O (5% to 10% v/v depending on frequency of spectrometer) was added to the sample. The parameters for the ^1^H-^15^N Heteronuclear Single-Quantum Correlation (HSQC) experiment are described in Supplementary Table 5.

### NMR analysis of the auxin co-receptor complex

In our system, the NMR experiments had to be conducted with 5-10 μM TIR1 protein at 4°C and were completed within 18 hours from finishing the purification. ^15^N isotopically-labelled AXR3 DI/DII protein and unlabelled TIR1 protein was prepared in a 1:3 ratio with 5% D_2_O and measured using a ^1^H-^15^N HSQC experiment following the parameters described in Supplementary Table 5. The full auxin co-receptor complex was studied by the addition of 200 μM auxin (unlabelled) to the sample. The NMR experiments were initiated with fresh TIR1 and completed within 18 hours of finishing the TIR1 purification.

### NMR backbone assignment

The following NMR experiments were used in the assignment of the backbone of AXR3 DI/DII: HNCA, HNcoCA, HNcaCB, CBcacoNH, HNcaCO, HNCO using ^13^C, ^15^N isotopically-labelled protein (290 μM). The parameters for are described in Supplementary Table 1. All the assignment experiments were performed at 600 MHz at 16.5°C using an Agilent DDX3 NMR spectrometer with a RT HCN triple resonance probe. The assignment data were analysed with minimal automation in the software CcpNmr Analysis.

### Sequential NMR backbone assignment through the prolines in the AXR3 DI/DII protein

A set of 2D ^13^CO detected NMR experiments CON, hCAnCO, and hCACO were used in the assignment of prolines in the carbon backbone of AXR3 DI/DII. The parameters for the NMR experiments are described in Supplementary Table 2. The experiments were performed at 950 MHz at 16.5°C using a TCI cryoprobe with a cooled amplifier on carbon.

### Identifying and estimating the occupancy of the *cis* and *trans* isomer populations

The ^13^C_α_ *cis* Pro population was predicted to have an up-field chemical shift of around 0.5 ppm^41,42^. The hCAnCO and hCACO spectra for AXR3 DI/DII show an up-field ^13^C_α_ chemical shift difference of around 0.3 ppm for the *cis* isomer population of P87 compared to the *trans* isomer position. The height and volume of NMR signals assigned to G85 and W86 were determined automatically from the assignment peak list for the ^1^H-^15^N HSQC spectrum within the software CcpNmr Analysis using the peak picking option. The height of the NMR signals was measured by a parabolic method. The NMR experiments were performed at least 12 hours after purification.

### ^15^N R_2_ relaxation of AXR3 DI/DII

A ^15^N R_2_ relaxation experiment was performed at 16.5°C and at 950 MHz following the parameters in Supplementary Table 6. Ten values of the R_2_ relaxation delay (S) were used, including two repeat values and recorded in a random order of 0.06, 0.39, 0.84, 0.26, 0.64, 0.13, 0.52, 0.26, 1.03, 0.64.

### Surface plasmon resonance

Streptavidin sensor chips (GE Healthcare Life Sciences) were used in all SPR assays and prepared with AXR2 degron peptide (IAA7: biot-AKAQVVGWPPVRNYRKN) and mutated AXR2 peptide (mIAA7: biot-AKAQVVEWSSGRNYRKN) as previously described^34^, using HBS-EP buffer (GE Healthcare Life Sciences) and a Biacore T200. TIR1 for these experiments was prepared as detailed above except that TIR1 protein was eluted with 100 μg ml^−1^ 3×FLAG peptide in 10 mM HEPES pH7.4, 150 mM NaCl, 3 mM EDTA, 1 mM TCEP, 0.05% Tween 20. Each SPR experiment consisted of at least 1 minute buffer baseline at a flow rate of 20-25 μL / minute followed by a 4 minute injection of 50 μM auxin with TIR1 protein in HBS-EP buffer (indicative protein concentration 1.75 μM, determined by UV absorbance at A_280_). A dose series of each auxin (IAA or cvxIAA) was injected over the AXR2 (IAA7) peptide^34^. A series of 8 auxin concentrations was run, with two being duplicated. A control with solvent (1% DMSO final, as for all auxin treatments) was used for a double subtraction baseline. All sensorgrams represent data for channel 2-1, with 4x mutated AXR2 (mIAA7) on channel 1^34^.

## Supporting information

Ramans Harborough et al 2019 Fuzzy Complex SI

## Acknowledgements

We thank Professor Sheena Radford FMedSci FRS for critical reading of the manuscript. We also thank Dr John Paul Evans and Dr Nathan Kidley for their support throughout the project. We acknowledge the NMR facility for access to the 950 MHz and 600 MHz spectrometers funded by the University of Leeds and the 750 MHz spectrometer funded by the Wellcome Trust (Award Reference: 094232). SPR was performed at the Wellcome Trust-supported Biomolecular Interactions facility in the Astbury Centre, Faculty of Biological Sciences, University of Leeds (Award Reference: 062164/Z/00/Z) and at the University of Warwick Synthetic Biology Research Centre (Award Reference: BB/M017982/1). This research was funded by grants from the Biotechnology and Biological Sciences Research Council (Award references: BB/L010623/1 to S.K. and R.N., and BB/I532402/1 to S.R.H.) and Syngenta UK (Award Reference: 1232512 to S.K. & S.R.H.). M.K.^3^ was supported by the EU MSCA-IF project CrysPINs (792329).

