## Supplementary material for "A fuzzy encounter complex precedes formation of the fully-engaged TIR1-Aux/IAA auxin co-receptor system": Ramans Harborough et al 2019 Fuzzy Complex SI

**Supplementary Information**

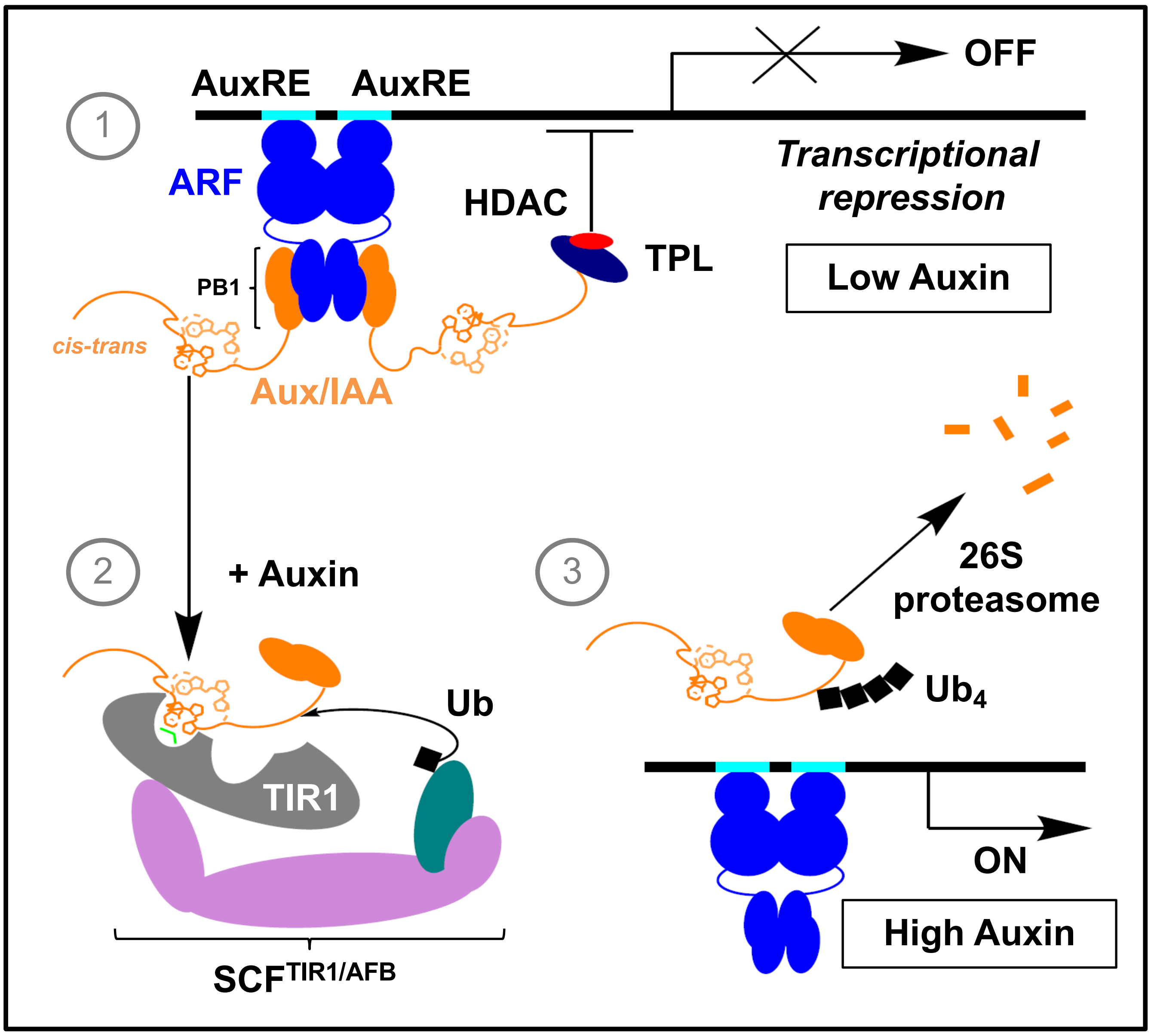


**Supplementary Figure 1 | Summary of the auxin signal transduction pathway.** Schematic summary of auxin signal transduction in the nucleus of a plant cell: (1) auxin response elements (AuxREs) in the promoters of auxin-inducible genes act as binding sites for the auxin response factor (ARF) transcription factors. ARF proteins, which themselves dimerize to bind pairs of AuxREs form further interactions with Aux/IAA transcriptional co-repressor proteins, via C-terminal PB1 interaction domains common to both families of proteins. The Aux/IAA recruits the co-repressor TOPLESS (TPL), which in turn may recruit histone deacetylase complexes (HDAC) to induce a transcriptionally repressed state at the targeted locus. (2) An increase in auxin levels promotes the formation of the auxin receptor complex. This is composed of an SCF E3 ubiquitin ligase ($SCF^{TIR1/AFB}$), where the F-box protein TRANSPORT INHIBITOR RESPONSE1 (TIR1) provides the binding sites for ligand and co-receptor Aux/IAAs (3) As a consequence, Aux/IAA proteins become polyubiquitinated, targeting them for rapid degradation in the 26S proteasome, and prompting the de-repression of auxin-regulated genes.

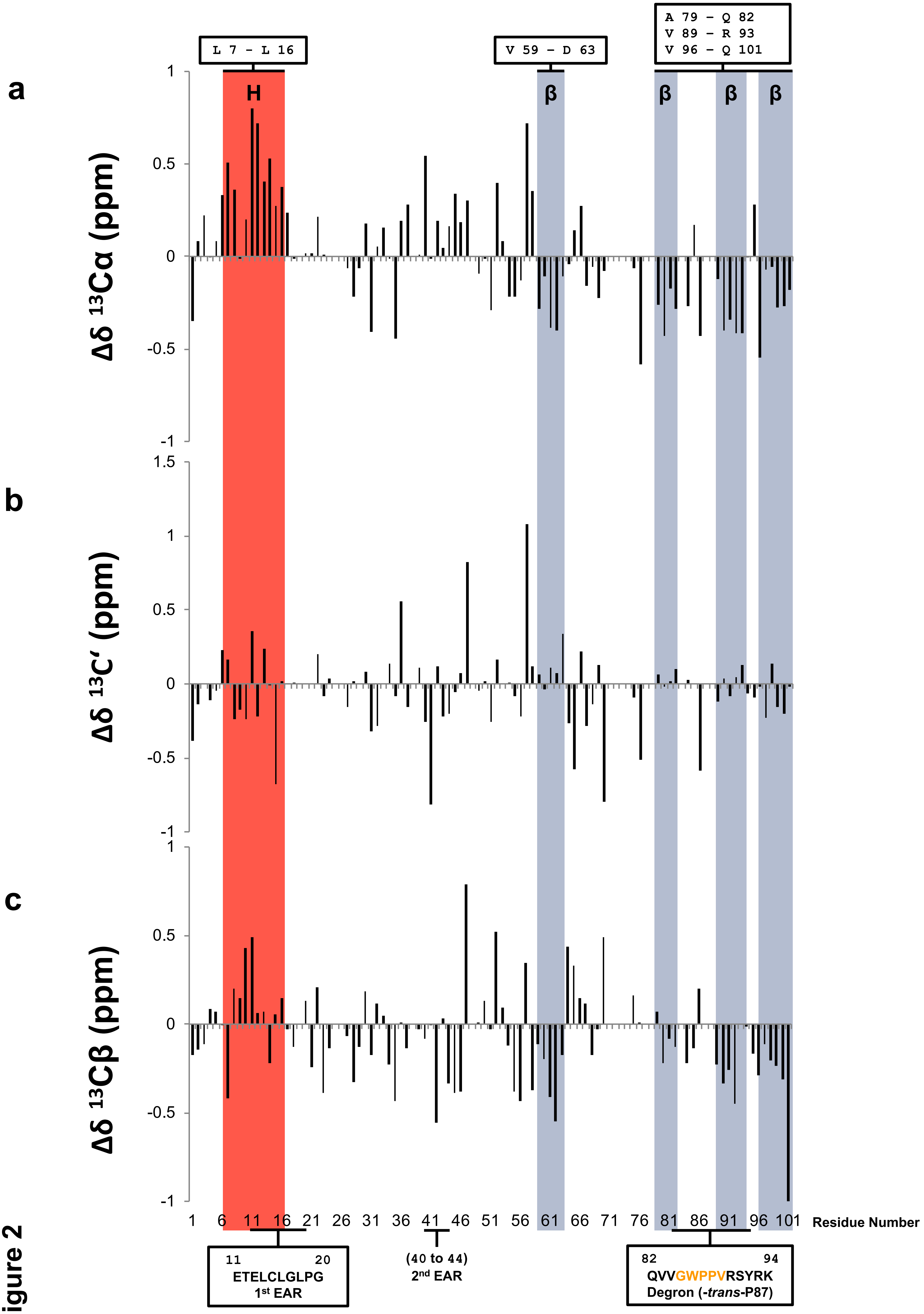


Supplementary Figure 2

**Supplementary Figure 2| Chemical shift indices for $^{13}C_{\alpha}$, $^{13}C'$, and $^{13}C_{\beta}$.** Positive chemical shift differences ($\Delta\delta$) for the $^{13}C_{\alpha}$, $^{13}C'$ indices indicate a tendency for helical secondary structure. Negative $\Delta\delta$ for the $^{13}C_{\alpha}$, $^{13}C'$ indices indicate a tendency for β-secondary structure. The opposite scenario applies to the $\Delta\delta$ $^{13}C_{\beta}$ index. Consensus regions for secondary structure tendencies between the $\Delta\delta$ $^{13}C_{\alpha}$, and $^{13}C'$ indices are highlighted light-red and dark-grey, for the helical and β-secondary structure respectively. Residues forming the termini of the consensus regions for secondary structure tendencies are box annotated. The degron and the EAR motifs are also included as reference points.

a

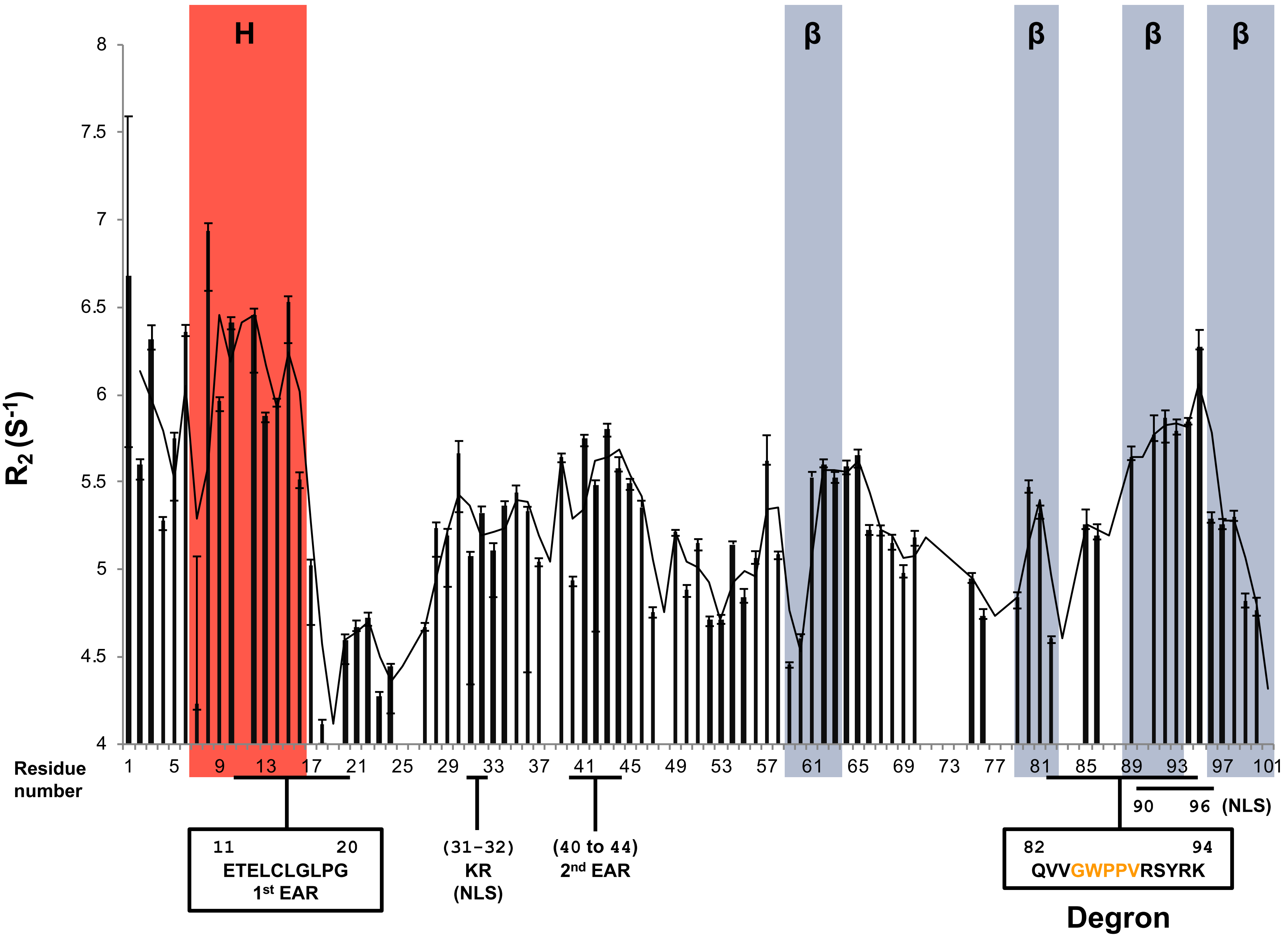


b

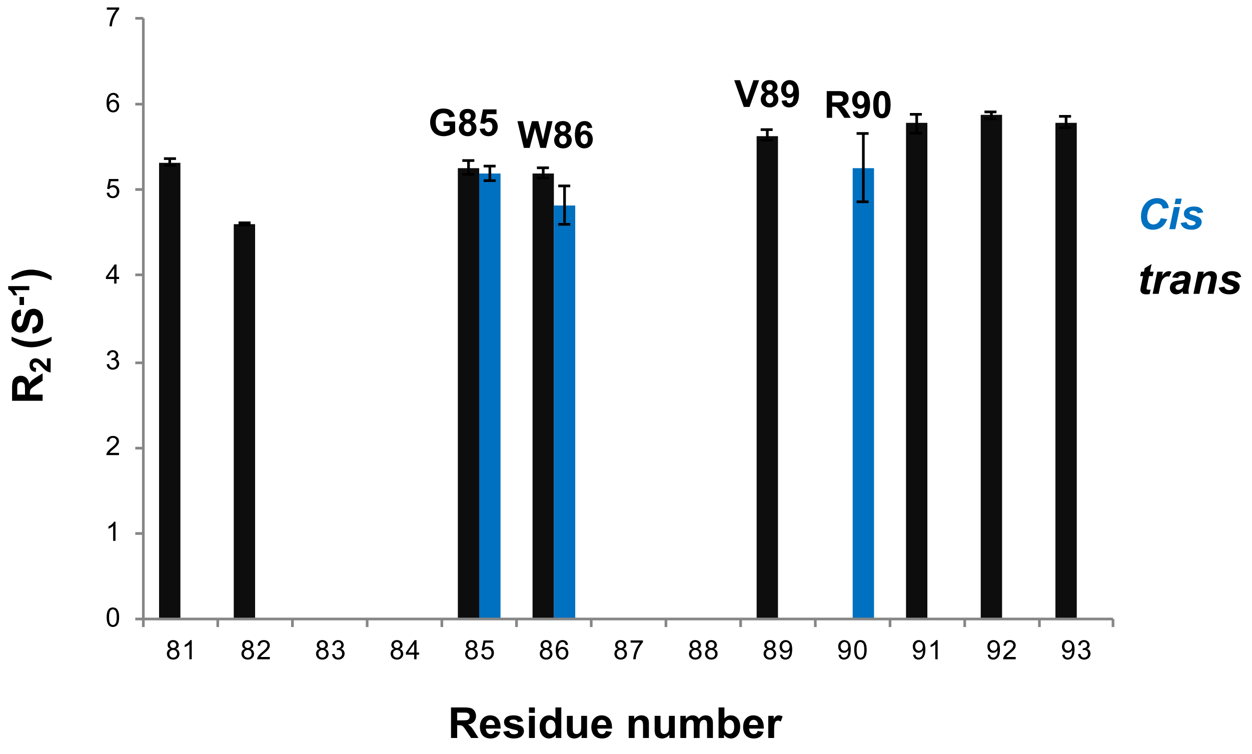


Supplementary Figure 3

**Supplementary Figure 3 | Comparison between $R_2$ relaxation rates and the consensus regions for secondary structure tendencies of AXR3 DI/DII.** (**a**) $R_2$ relaxation rate for residues along the carbon backbone of AXR3 DI/DII. Relative increases in the $R_2$ rate are associated with regions of reduced flexibility. The $R_2$ data set is overlaid with consensus regions for secondary structure tendencies as indicated by the chemical shift indices. Helical regions are shown in light-red and β-secondary structure shown in dark-grey. The data set is annotated with consensus sequences for functional domains I and II, including the nuclear localisation signal (NLS) KR motif which functions in partnership with a 2$^{nd}$ NLS at R90 to V96. Gaps in the $R_2$ data set are associated with prolines and residues in signal dense regions preventing peak picking. (**b**) $R_2$ relaxation rate for residues in domain II. Showing the $R_2$ rates for the split NMR resonances for degron residues G85 to R90 within the $^{1}H$-$^{15}N$ HSQC spectrum that are linked to the *cis* (blue) and *trans* (black) isomers of P87.

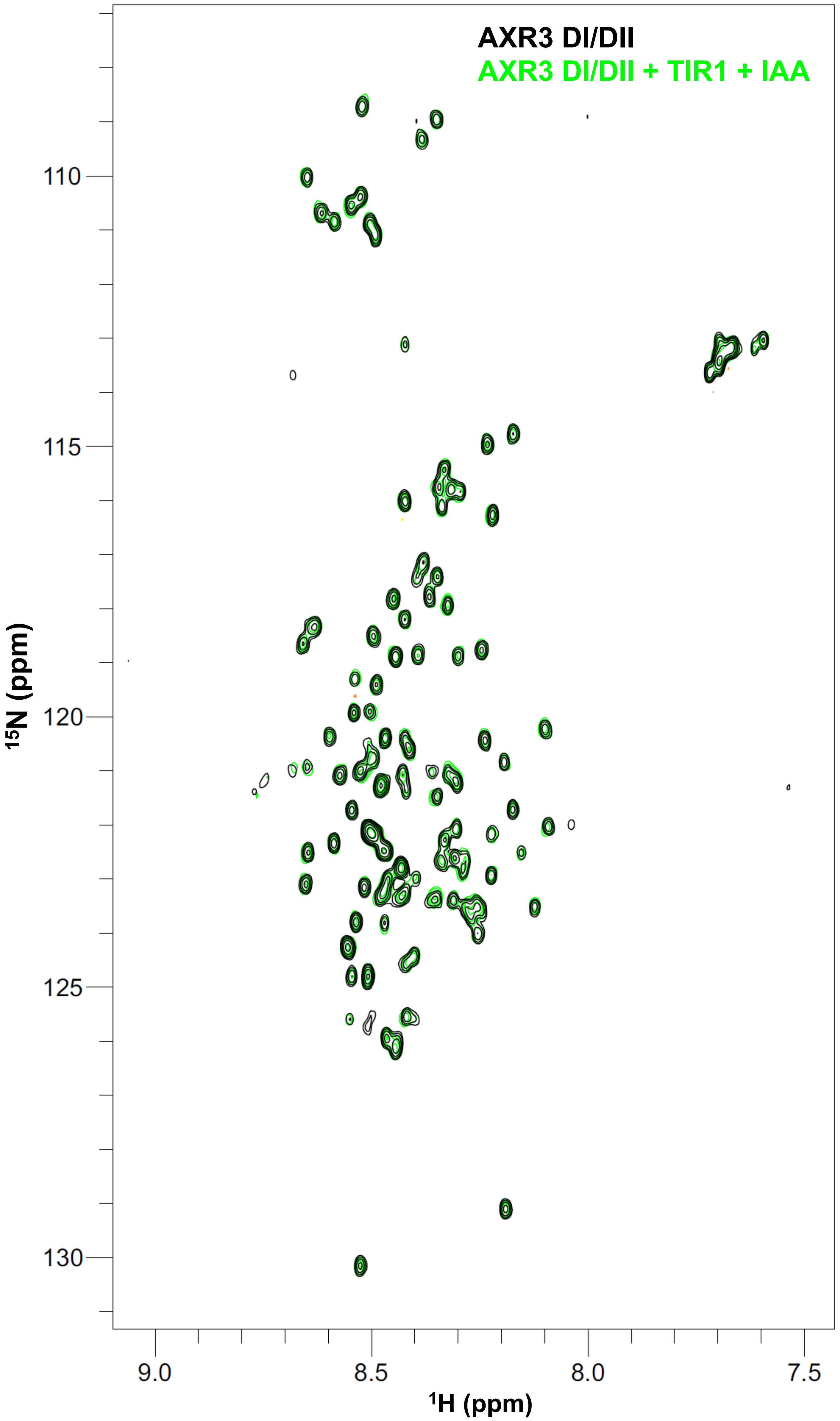


**Supplementary Figure 4 | AXR3 DI/DII in complex with auxin and TIR1 retains its intrinsic disorder.** $^{1}H$-$^{15}N$ HSQC spectrum of the isotopically labelled ($^{15}N$) protein AXR3 DI/DII at 4 °C both alone (black), and with unlabelled TIR1 in the presence of the unlabelled auxin (IAA) (green).

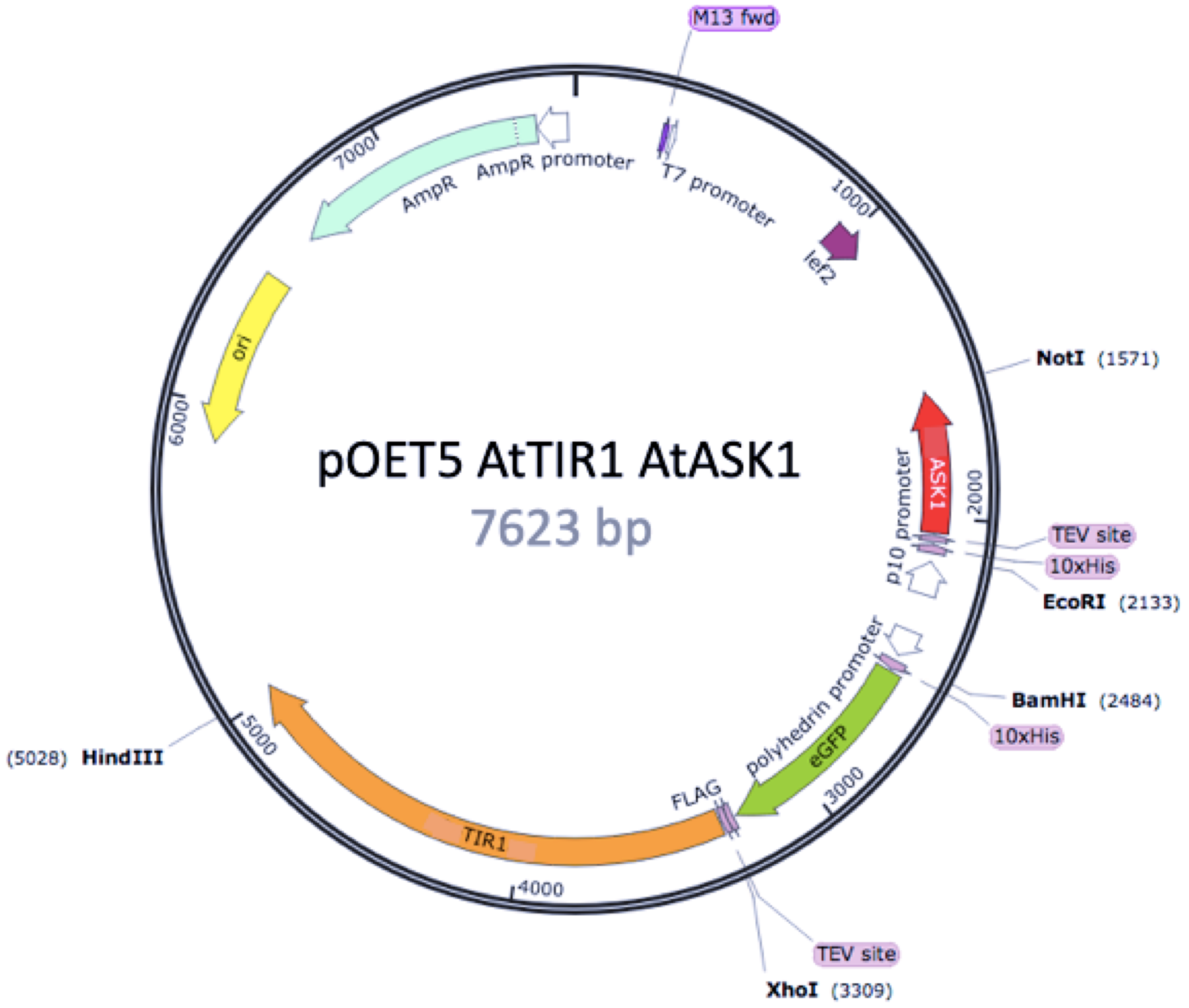


**Supplementary Figure 5 | Map of baculovirus transfer vector for co-expression of AtTIR1 and AtASK1.** Codon-optimized gBlocks (Integrated DNA Technologies) for His10-eGFP-FLAG-(TEV)-AtTIR1 and His10-(TEV)-PpASK1 were cloned into the pOET5 transfer vector (Oxford Expression Technologies). The map was created using SnapGene® software (from GSL Biotech; available at snapgene.com).

**Supplementary Table 1. Parameters for assignment experiments in the analysis of $^{13}C$ and $^{15}N$ isotopically labelled AXR3 DI/DII protein.** All experiments were performed at 16.5˚C, 600 MHz and with recycling delays of 1 second. Four scans were collected.

| Experiment | Nuclei | | | Spectral width (Hz) | | | Number of complex data points | | |
|---|---|---|---|---|---|---|---|---|---|
| | t1 | t2 | t3 | t1 | t2 | t3 | t1 | t2 | t3 |
| HNCA | $^{15}N$ | $^{13}C$ | $^{1}H$ | 1600.0 | 4525.3 | 6613.8 | 32 | 64 | 1024 |
| HNcoCA | $^{15}N$ | $^{13}C$ | $^{1}H$ | 1600.0 | 4525.3 | 6613.8 | 32 | 64 | 1024 |
| HNcaCB | $^{15}N$ | $^{13}C$ | $^{1}H$ | 1600.0 | 10558.9 | 6613.8 | 32 | 64 | 1024 |
| CBcacoNH | $^{15}N$ | $^{13}C$ | $^{1}H$ | 1600.0 | 12067.3 | 6613.8 | 32 | 64 | 1024 |
| HNcaCO | $^{15}N$ | $^{13}C$ | $^{1}H$ | 1600.0 | 1600.0 | 6613.8 | 48 | 64 | 1024 |
| HNCO | $^{15}N$ | $^{13}C$ | $^{1}H$ | 2500.0 | 1600.0 | 9615.4 | 48 | 64 | 1024 |

**Supplementary Table 2. Parameters for proline assignment experiments in the analysis of $^{13}C$, $^{15}N$ isotopically labelled AXR3 DI/DII protein.** All experiments were performed at 16.5˚C, 950 MHz with recycling delays of 1.5 seconds and acquisition time of 71.3 ms.

| Experiment | Scans | Nuclei | | Spectral width (Hz) | | Number of complex data points | |
|---|---|---|---|---|---|---|---|
| | | t1 | t2 | t1 | t2 | t1 | t2 |
| CON | 8 | $^{15}N$ | $^{13}C$ | 3851.5 | 7183.9 | 160 | 512 |
| hCACO | 8 | $^{13}C$ | $^{13}C$ | 7168.5 | 7183.9 | 64 | 512 |
| hCAnCO | 16 | $^{13}C$ | $^{13}C$ | 7168.5 | 7183.9 | 180 | 512 |

**Supplementary Table 3. Steady state affinity parameters.** Kinetic parameters calculated in BiaEvaluation software (GE Healthcare) using the 1:1 Langmuir model

| Degron | $K_D$ (µM) | $R_{max}$ | Offset (RU) | $Chi^2$ ($RU^2$) |
|---|---|---|---|---|
| IAA | 2.75 | 550 | 9.93 | 227.00 |
| cvxIAA | 26.40 | 182 | -0.51 | 8.91 |

**Supplementary Table 4. Primer list.**

| Experiment | Primer | Sequence |
|---|---|---|
| Cloning AXR3 DI/DII | B1IAA17 | 5'- GGGGACAAGTTTGTACAAAAAAGCAGGCTGCATGATGGGCAGTGTCGAGCTGAA TCT -3' |
| | IAA17d stopB2 | 5'- GGGGACCACTTTGTACAAGAAAGCTGGGTATCATTTTTGGCAGGAAACCATCAC G -3' |
| Site-Directed mutagenesis: *axr3-3* | IAA17SD MF33 | 5'- TGGCCACCGGGGAGATCATACCGGAAGA -3' |
| | IAA17SD MR1 | 5'- TCCCACAACTTGTGCCTTGGCCGGAGGT -3' |

**Supplementary Table 5. Parameters for heteronuclear single-quantum correlation (HSQC) experiment.** HSQC experiments to study the auxin co-receptor complex were performed at 4˚C and 950 MHz. All other HSQC experiments were performed at 16.5 ˚C and either 950 or 750 MHz.

| **Experiment** | **Recycling Delays (S)** | **Scans** | **Nuclei** | | **Spectral width (Hz)** | | **Number of complex points** | |
|---|---|---|---|---|---|---|---|---|
| | | | t1 | t2 | t1 | t2 | t1 | t2 |
| HSQC (co-receptor complex study) | 1 | 16 | $^{15}N$ | $^{1}H$ | 2407.2 | 15243.9 | 90 | 1024 |

**Supplementary Table 6. $^{15}N$ $R_2$ relaxation experiment of AXR3 DI/DII protein.** All experiments were performed at 16.5˚C and 950 MHz.

| **Experiment** | **Recycling delay (S)** | **$R_2$ Recycling delay (S)** | **Scans** | **Nuclei** | | **Spectral width (Hz)** | | **Number of complex data points** | |
|---|---|---|---|---|---|---|---|---|---|
| | | | | t1 | t2 | t1 | t2 | t1 | t2 |
| $R_2$ relaxation | 2.8 | 0.01612 x L* | 4 | $^{15}N$ | $^{1}H$ | 2599.7 | 15243.9 | 200 | 1024 |

* where L value was changed after each run in the following sequence:4, 24, 52, 16, 40, 8, 32,16, 64, 40. Giving relaxation delay values of 0.06, 0.39, 0.84, 0.26, 0.64, 0.13, 0.52, 0.26, 1.03, 0.64.
